# Novel two-component system-like elements reveal functional domains associated with restriction-modification systems and paraMORC ATPases in bacteria

**DOI:** 10.1101/2020.08.26.268516

**Authors:** Daniel Bellieny-Rabelo, Willem JS Pretorius, Lucy N Moleleki

**Author notes:** Corresponding author. (LNM). Department of Ecology and Environmental Sciences, Umeå University, Umeå, 90736, Sweden (Daniel Bellieny-Rabelo).

## Abstract

**Background:** Two-component systems (TCS) are essential machineries allowing for efficient signal recognition and transmission in bacterial cells. The majority of TCSs utilized by bacteria are composed by a sensor histidine kinase (HK) and a cognate response regulator (RR). In the present study, we report two newly predicted protein domains — Response_reg_2 (PF19192) and HEF_HK (PF19191) — in bacteria which exhibit high structural similarity, respectively, with typical domains of RRs and HKs.

**Results:** Additionally, the genes encoding for the novel predicted domains exhibit a 91.6% linkage observed across 644 genomic regions recovered from 628 different bacterial strains. The remarkable adjacent co-localization between genes carrying Response_reg_2 and HEF_HK in addition to their conserved structural features, which are highly similar to those from well-known HKs and RRs, raises the possibility of Response_reg_2 and HEF_HK integrating a new TCS in bacteria. The genomic regions in which these predicted two-component systems-like are located additionally exhibit an overrepresented presence of restriction-modification (R-M) systems especially the type II R-M. Among these, there is a conspicuous presence of C-5 cytosine-specific DNA methylases which may indicate a functional association with the newly discovered domains. The solid presence of R-M systems and the presence of the GHKL family domain HATPase_c_3 across most of the HEF_HK-containing genes are also indicative that these genes are evolutionarily related to the paraMORC family of ATPases.

**Conclusions:** The present study uncovered two novel protein conserved domains and raised the possibility of a TCS-like system undertaking a regulatory role mechanistically linked to R-M systems in a large variety of bacterial lineages.

## INTRODUCTION

Monitoring environmental conditions is determinant for the success of microbial organisms. To address this demand, biochemical cascades that enable programmatic responses to diverse stimuli have evolved across the three domains of life [1,2]. A dominant signaling archetype in nature is comprised by a receptor/transmitter protein paired with a ligand-binding regulator to be affected downstream, collectively known as two-<u>c</u>omponent <u>s</u>ystems (TCS) [3,4]. Specifically in bacteria, preferable conservation of <u>h</u>istidine-<u>k</u>inase (HK) as the transmitter module in receptor proteins has been recurrently documented since the discovery of TCSs roughly 3 decades ago [5-8]. HKs commonly associate with intracellular response regulators (RR), which transduce signals and generate responses through diverse mechanisms, which may range from binding DNA or proteins, to enzymatic catalysis [9-11]. As the genomic TCS collection in a single cell may comprise dozens of duplets, interaction specificity amongst the cognate partners of a TCS is critical in order to avoid cross-talk with other pathways [12,13].

In this context, the canonical transduction chain in this system depends on five key domains. The first domain is a highly conserved <u>c</u>atalytic <u>A</u>TP-binding domain (CA) typified by the HATPase domains (Pfam: PF02518; PF13581; PF13589; PF14501) which integrates the C-terminal regions of all known HKs and harbors a characteristic nucleotide-binding α/β sandwich known as the Bergerat fold [14,15]. The proteins harboring this structural fold are classified into the GHKL superfamily [15]. The second domain is a <u>d</u>imerization domain containing the active <u>H</u>is site for <u>p</u>hosphorylation (DHp). Together, DHp and CA domains comprise the two well conserved regions in HK proteins. The prototypic DHp domain harbors the active site His in a double α-helical structure termed H-box [16]. Structural variants of DHp have been described over the years, which are commonly called HiskA domains (Pfam: PF00512; PF07568; PF07730; PF07536) [16-18]. The third well conserved domain in TCS is the <u>rec</u>eiver domain (REC), which is commonly found in the RRs, and features a highly conserved Asp residue within its active site (Pfam: PF00072) [19]. This active site Asp is typically found within a characteristic (βα)_5_ fold conserved in RR proteins [20]. The remaining two domains among the aforementioned five typical domains in TCS are highly variable and comprise: (a) a sensor/input domain (in HKs) and (b) effector/output (in RRs), respectively responsible for recognizing specific signals, and eliciting responses. The high sequence variability observed in these domains enables a myriad of signal/target recognition patterns [11].

Functional classification of HKs usually provides necessary insight that allows initial characterization of the TCS role. This classification is generally inferred by means of domain architecture, which encompasses trans<u>m</u>embrane regions (TMRs) and sensor domain arrangement within the sequences [21]. Three major groups of HKs can be determined following this criterion. The largest amongst these groups is represented by the periplasmic-sensing HKs carrying an extracellular sensor typically flanked by two TMRs [21]. Members of this group commonly harbor a so-called ‘linker ‘domain (e.g. HAMP, or PAS) flanked by the TMR2 and the DHp domain in the protein sequence [22]. The second major group include both soluble and membrane anchored sensors typified by inward facing input domains [21]. Among these, NtrB along with its cognate NtrC comprise a well-characterized TCS model responsible for gene regulation under nitrogen depravation scenarios in *Escherichia coli* [5,23]. The role of another soluble HK termed HoxJ has also been established in *Ralstonia eutropha* as being part of a regulation mechanism for hydrogenase genes in response to H_2_ [24,25]. Interestingly, *hoxJ* is invariably found adjacently to *hoxBC* genes, which encode two proteins that helps assemble the sensory/transduction complex HoxJ-BC necessary for proper H_2_ perception [21]. This is unsurprising given the typical co-localization of functionally related genes in the same gene-neighborhoods found in prokaryotic genomes [26]. HKs of the third group are characterized by the conservation of 2-20 TMRs that actively enable stimulus perception often associated with the membrane itself (e.g. mechanical stress) [21,27].

In this report, we predict two novel domains structurally related respectively to DHp and REC in bacteria. In addition, the extensive co-localization observed in genes that encode these domains support the possibility of these domains comprising a novel TCS. These genes also exhibit a tendency of being neighbored by restriction-modification systems, which provided further insights on their putative biological role.

## RESULTS

### Establishing homologous sets and inspecting genomic contexts

On the course of a large-scale gene expression analysis conducted by our group on potato tubers using the highly virulent plant pathogen *Pectobacterium brasiliense* strain PBR 1692 (*Pb*1692), three adjacent uncharacterized genes were detected exhibiting significant activation [28,29]. One of these three genes (*PCBA_RS21900*) was identified as a member of the GHKL superfamily. The encoded product of *PCBA_RS21900* exhibits 2 HATPase domains (Pfam: PF02518; PF13589) [15]. These domains are typically found in Hsp90 chaperones, DNA-gyrases, MutL and HK family members [15]. The other two neighboring genes (*PCBA_RS05725-05720*), contrarily, did not harbor any known conserved domains. With further inspection of the *PCBA_RS05725-05720* homologs in other strains from the same organism (i.e. LMG21371, PcbHPI01) through nucleotide alignment, we found that these two hypothetical entries probably comprise in fact one single gene. Curiously, these homologs from LMG21371 and PcbHPI01 were also neighbored by GHKL superfamily members. Hence, by aiming to improve the accuracy in the subsequent computational analyses involving sequence comparisons, the sequence from *Pb* LMG21731 (*KS44_RS10365*) was used instead of those from *Pb*1692. The goal here was to provide an assessment on the frequency of genomic co-localization between *KS44_RS10365*-related genes and GHKL superfamily members in other bacterial genomes.

In order to perform an initial gene-neighborhood screening on KS44_RS10365-related sequences, we collected a wide range of similar protein sequences from prokaryotes in an extensive search on the NCBI database (see ‘Methods ‘for details). From the initial sequence search, 677 bacterial strains returned positive hits for KS44_RS10365, and their respective genomic data were obtained to conduct subsequent analyses [TableS1]. Next, in order to predict orthology relationship among the 441,897 sequences collected from the 677 strains, a sequence clustering step was conducted using OrthoMCL [30]. The sequence clustering analysis produced 23,397 distinct <u>o</u>rthologous <u>g</u>roups (OG) in total, which were populated by ranges of 2-990 sequences. This approach was paralleled by conserved domains inspection [31] in the same protein data set. Conserved domains assessment was used to provide functional annotations, and to carefully check the cohesiveness of domain architectures within the clustered sequences in the OGs.

The integration of domains and OGs annotation with genomic coordinates enabled extensive gene-neighborhood analysis on the genes encoding the OG_4 members, which harbors KS44_RS10365 and its predicted orthologs. The OG_4 sequences spread through 628 different strains in which 30 conserve 2-3 paralogs, totaling 664 proteins [TableS2]. Out of these 664 paralogs, 20 were in regions unsuitable for contextual analysis due to incompletion of the respective genome assemblies, which resulted in the lack of necessary data in the region of interest. In the 644 analyzed regions (after the exclusion of the 20 unsuitable regions), we observed a remarkable linkage of <u>91.6%</u> between OG_4 and GHKL superfamily members (OG_5), in a distance of up to three genes. Next, by scanning a broader region including 10 genes up/downstream to OG_4 homologs, the HATPase-containing genes (GHKL) were among the most prevalently detected domains (Figure 1). In the OG_4 members genomic neighborhoods we also found the dominant presence of DNA_methylase (Pfam: PF00145) containing genes, frequently located upstream of the OG_4 homologs. This strong co-localization raised the possibility of the association of OG_4 members with restriction-<u>m</u>odification (R-M) systems, which will be addressed in detail in a subsequent topic. Other domains associated with two-component signal transduction systems, such as HiskA (Pfam: PF00512) and Response_reg (Pfam: PF00072) were also frequent in the OG_4 members regions (Figure 1). The presence of immunity- and polymorphic toxin-related domains such as CdiI_3 (Pfam: PF18616) and Ntox28 (Pfam: PF15605) in OG_4 members ‘vicinity indicates that these genes can frequently be found in association with toxin-antitoxin systems. Additionally, several genes located in these regions for which no known domains could be detected were grouped together in orthologous groups, such as the members of OG_20, OG_16, OG_24, OG_8 and OG_294 (Figure 1). The traceable evolutionary relationship among these uncharacterized genes can potentially drive the discovery of novel functionalities in subsequent studies. Together, these observations revealed a strong co-localization pattern among OG_4 and OG_5 (GHKL-containing) members in a wide range of bacteria as well as the most important functional groups to which they frequently associate with. Below, we lay emphasis on sequence analysis in order to examine other conserved features in OG_4/5 groups aiming to determine which biological role they might undertake.

**Figure 1.**
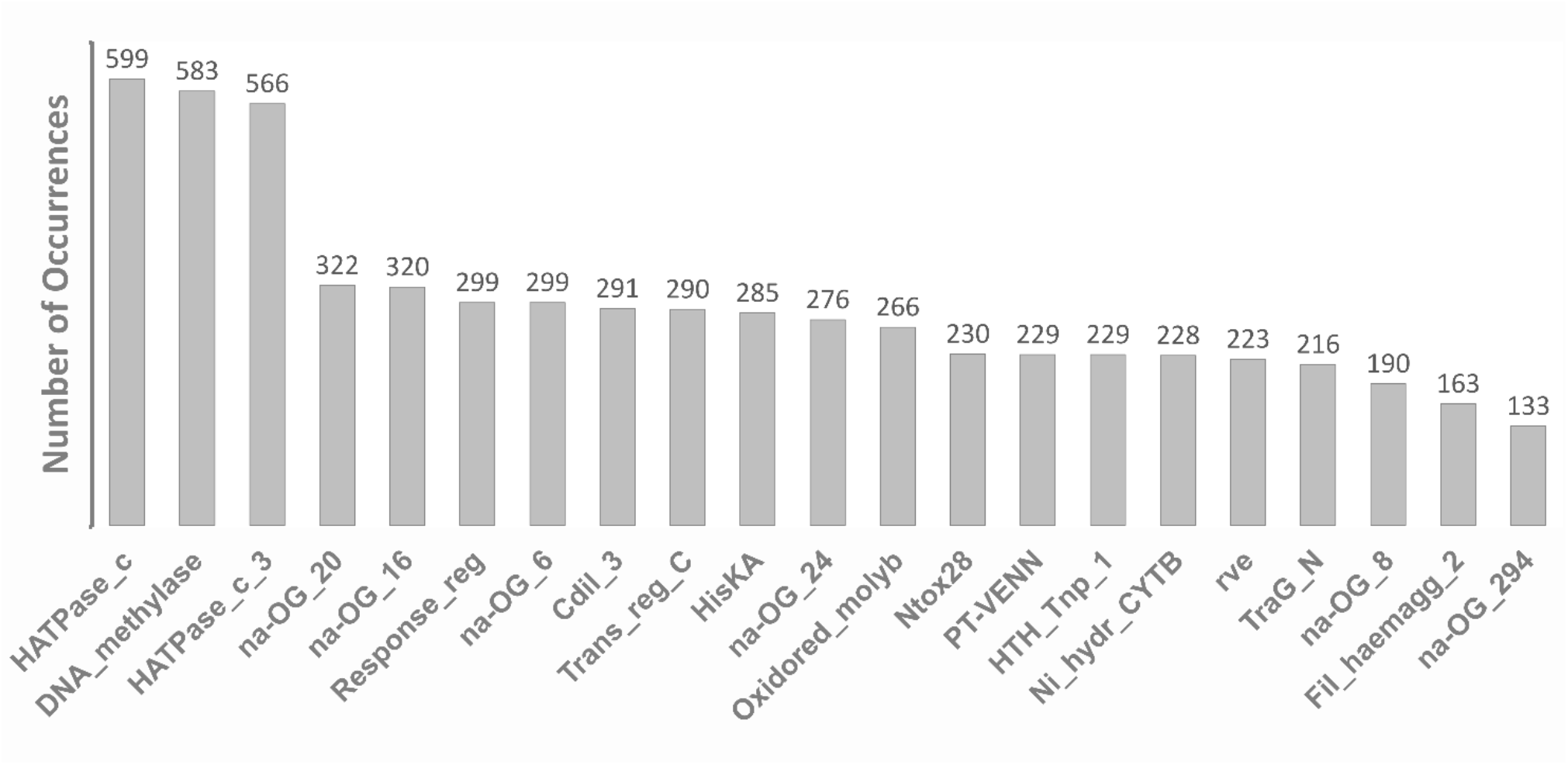
Top 1% best represented protein domains encoded in OG_4 members’ genomic regions. 628 genomes containing OG_4 genes were analyzed 10 genes up- and downstream. The functional domains encoded by each loci in that range were computed and the total amounts are displayed in the bar plot. Domain names were detected by hmmscan tool and are annotated according to the Pfam database. For each gene product, in the absence of known domains (na – not applicable), the respective orthologous group label (prefix: OG_; suffix: numeric tag) are displayed.

### Domain architectures characterization

As a preliminary step, we performed a sequence analysis step using the OG_5 sequences in comparison with the major GHKL families (i.e. Hsp90, DNA-gyrases, MutL and HK). This revealed that OG_5 sequences only produced results when aligned with HKs. This preliminary observation, in combination with the remarkable trend for co-localization of OG_4 and 5 indicated that OG_5 and OG_4 could possibly comprise a TCS. Hence, taking advantage of diverse REC and DHp domain profiles publicly available [32], members of OG_4 and OG_5 were analyzed along with those profiles in order to verify the possible presence of similar features in the <u>m</u>ultiple-<u>s</u>equences <u>a</u>lignments (MSA). These analyses revealed two previously unknown conserved regions respectively in OG_4 and OG_5 bearing strong similarities with REC and DHp domains.

The most pronounced feature of REC domains is the active site Asp residue [20], which was remarkably conserved with 100% consensus found across representative OG_4 sequences (Figure 2A). The active site Asp was also consistently conserved within the C-termini ends of β_3_. It has also been reported that two consecutive negatively charged residues located N-terminal to the active site are involved in co-factor coordination [33]. These residues were strikingly conserved in the C-terminal end of β_1_ both in OG_4 and in the canonical REC structure (Figure 2A and B). The Thr column located at the C-termini of β_4_ comprises another residue known to be conserved in REC domains, which was also shown to be highly conserved in the OG_4 representatives exhibiting high consensus level within this group. Additionally, the C-terminal end of β_5_ has a conserved Lys residue in the canonical REC, which was also observed in the OG_4 MSA. Although there was a lowly supported split of β_5_ structure into 2 β sheets, here, the C-terminal end of the second sheet seems highly preserved in OG_4 sequences exhibiting strong conservation of the Lys residue similarly to the REC domain (Figure 2A). This same β_5_ split probably reflects the difficulty in secondary structure prediction for some sequences in the MSA, which is not frequently observed when some of the sequences are inspected individually (Figure 2B). Notably, the tolerance in OG_4 sequences towards loop extensions interspersing the β/α structures, specially between α_1_/β_1_ and α_5_/β_5_, have presumably deepened the evolutionary distance to the canonical Response_reg (REC domain) resulting in the solid evolutionary separation between these motifs (Figure 2A and C). Although the OG_4 conserved motif and Response_reg have clearly diverged, the topology found in these domains exhibit remarkable similarity in the characteristic (βα)_5_ regulatory fold. This observed (βα)_5_ also exhibit key amino acid residues in identical positions to those reported in the canonical domain structure of REC domains.

**Figure 2.**
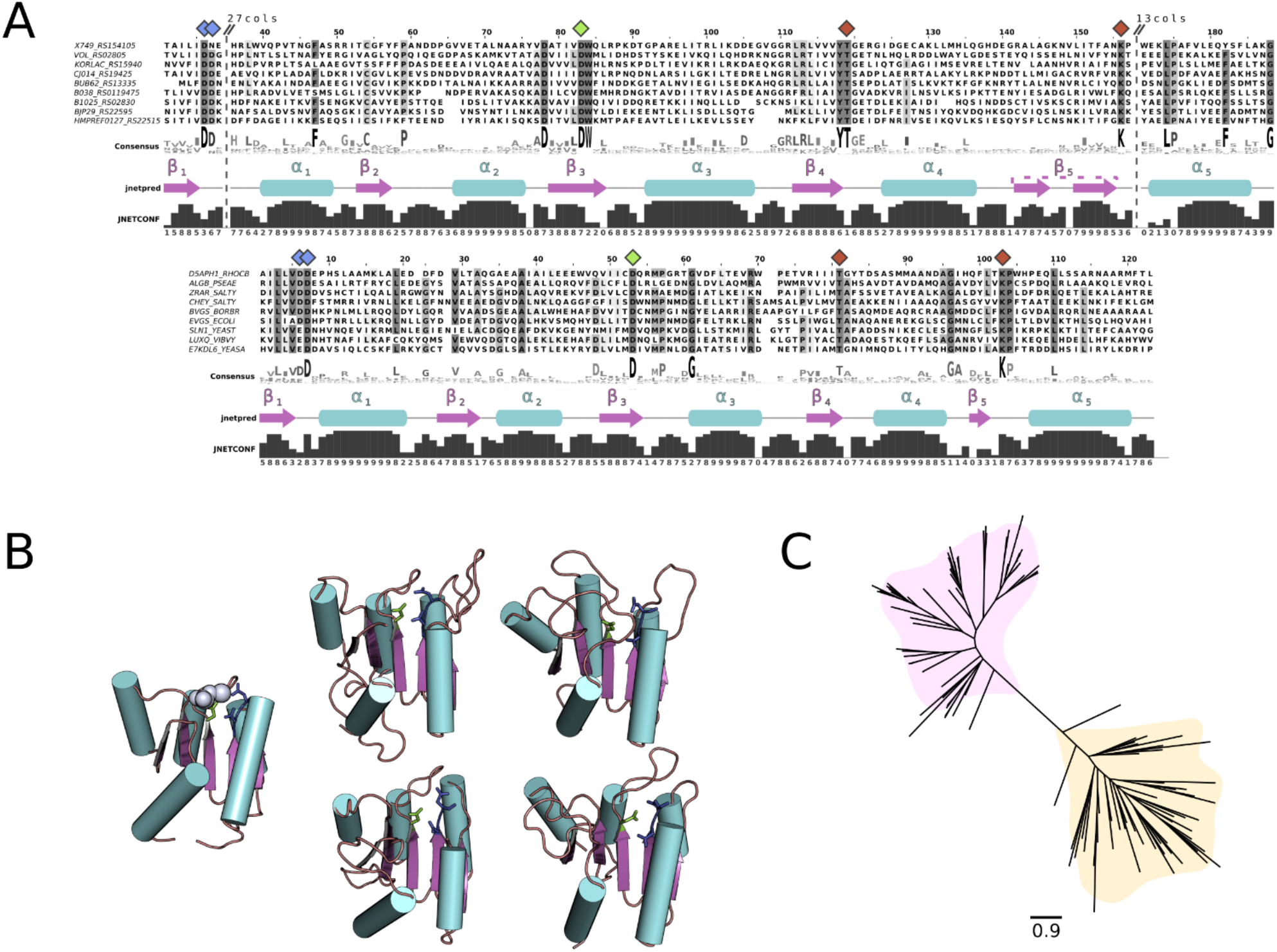
Multiple sequence alignment, structure scaffold and phylogenetic reconstruction of a conserved motif found in OG_4 members. **(A)** Multiple sequences alignments depicting the conserved regions in the REC domains in representative OG_4 (top) and Response_reg-containing (bottom) (Pfam: PF00072) proteins. Dashed lines in the alignments represent non-conserved loops hidden from the alignment. The numbers on the top indicate the number of amino acids not shown. Diamond shapes on top of specific columns represent key amino acids in the typical REC domain structure. Corresponding colors of diamond shapes in the top and bottom alignments represent homologous amino acids in both OG_4 and Response_reg-containing sequences. Numbered secondary structure (ss) predicted by JPred are respectively represented below each alignment. Confidence levels of ss predictions provided by JPred are displayed below the graphical ss representations. **(B)** Structures predicted by RaptorX represented as cartoons depict the canonical REC domain from CheY (PDB ID: 2FKA) with the Mg^2+^ highlighted in light purple on the left, and four examples of the newly described DHp domains from OG_4 representatives on the right: BJP29_RS22595 from *Bacteroides fragilis*; H147_RS0115650 from *Loktanella vestfoldensis*; B038_RS0119475 from *Martelella mediterranea*; VOL_RS02805 from *Vibrio owensii*. Three key Asp amino acids known to play a role in Mg^2+^ coordination are highlighted in the cartoons using the same color patterns as in the correspondent diamond shapes in (A). **(C)** Phylogenetic reconstruction of the Response_reg domains (yellow) and the predicted domains in OG_4 sequences (pink) is represented as an unrooted radial tree.

On the other hand, domain alignments of OG_5 members suggested a close relatedness between a conserved region within these sequences and the HiskA domain (Pfam: PF00512). Thus, by framing OG_5 and HiskA-containing representative sequences, the remarkable conservation of core His residue followed by a negatively charged residue (Glu or Asp) was observed (Figure 3A). The two predicted α-helices in the analysis comprised the characteristic H-box found in DHp, in which α_1_ harbored the core His residue (Figure 3A and B). Four randomly picked models of OG_5 sequences depicted the presence of the core His residue in α_1_ (Figure 3B). Such strong conservation of H-box helices constituted an important functional evidence, since this structure is responsible for both dimerization among HKs and phosphorylation of the active site Asp residue in the cognate RR. The phylogenetic reconstruction of H boxes from representative sequences of four canonical DHp domain families (see ‘Methods ‘for details) suggested that the domain found in OG_5 sequences might have evolved from HiskA (Figure 3C). These two closely related domains observably conserved a negatively charged residue (Glu or Asp) adjacent to the core His (Figure 3A). Since HKs attachment to the membrane, or lack thereof, is intrinsically associated with their function, we quantitatively compiled trans<u>m</u>embrane region (TMR) predictions on representative sequences carrying those same four DHp domains (i.e. HiskA, HiskA_2, HiskA_3 and HWE_HK). The results revealed a strong preference by HiskA_3 representatives to conserve 6 TMRs, whereas most of HWE_HK, HiskA_2 and OG_5 representative members tend to be found in soluble HKs (Figure 3D). In OG_5 proteins specifically, none of the sequences exhibited positive TMR predictions, which suggests that these group members function as soluble proteins. Taken together, these results strongly suggest that members of OG_4/5 groups could comprise a cytoplasmic TCSs. The two respective novel domains found in OG_4 and OG_5 will be hereafter referred to as Response_reg_2 and HEF_HK, and the sequences carrying these domains RR-like and HK-like respectively.

**Figure 3.**
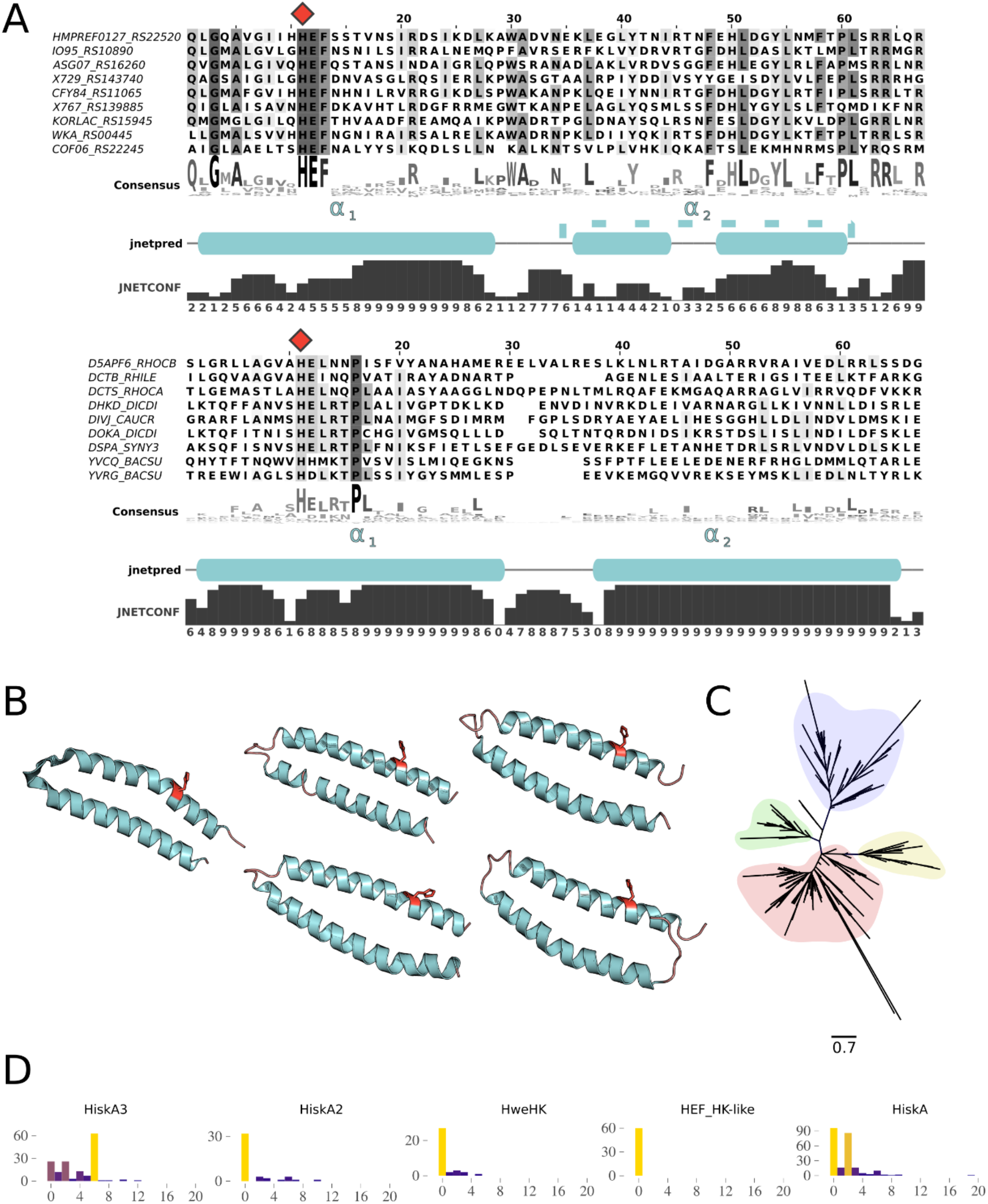
Multiple sequence alignment, structure scaffold and phylogenetic reconstruction of a conserved motif found in OG_5 members. The MSA representation follows the same pattern as in Fig. 1. **(A)** Red diamond shapes on top of MSA indicate the core His residue in both groups. **(B)** Structural scaffolds represent the α-helices of the canonical HiskA domain (left) (PDB_ID: 5B1N) and of four representatives of the newly predicted DHp domain (right): CFY84_RS11065 from *Acinetobacter apis*; HMPREF0127_RS22520 from *Bacteroides* sp. 1_1_30; WKA_RS00445 from *Escherichia coli*; KORLAC_RS15945 from *Kordiimonas lacus*. The core His residue in all five scaffolds are highlighted in red stick representations. **(C)** Phylogenetic reconstruction of HiskA variants, including HiskA (red), HiskA2 (green), HWE_HK (green), HiskA3 (blue), and the predicted DHp domain (yellow). **(D)** Each histogram depicts the frequency of transmembrane occurrences distribution found in representative sequences from each domain family predicted by TopCons.

### The association of RR-like and HK-like with R-M systems

Next, in order to gain further insight on the members of both families through “guilty by association” inference, we conducted in-depth genomic context analysis in these RR-like-harboring regions. By using the previously obtained 644 genomic regions from 628 bacterial strains carrying RR-like orthologs as reference, we first examined the functional domains encoded by their neighboring genes. This step aimed to inspect a genomic range of 10 neighboring genes up and downstream of the RR-like members. In addition, based on the previous assessment showing the solid presence of C-5 cytosine-specific DNA methylase (C5 Mtase) in the RR-like vicinity, we laid special emphasis on identifying R-M elements. To keep this analysis stringent from the functional prediction perspective (see ‘Methods ‘for details), we used the ‘gold standard ‘library of R-M-associated proteins from the REBASE database [34]. The REBASE gold standard set comprises only experimentally characterized components which have been associated with either restriction or modification functions. Aiming to determine which known functional domains are associated with R-M systems, we scanned the entire REBASE gold standard protein set using the Pfam database as reference. Subsequently, by inspecting the domains encoded by the genes neighboring the RR-like encoding genes and comparing these with the previous analysis, we found a consistent presence of R-M-related domains.

Next, in order to evaluate the correlation of R-M systems elements in close proximity to RR-like genes, a computational simulation step was implemented. This technique shuffles the entire gene pools from known genomes, generating ‘pseudo-genomes’. By analyzing the results from these randomized pseudo-genomes, it was possible to infer up to which frequency particular gene associations might occur by chance, as well as overrepresented associations. Here, we utilized 25 randomly picked complete bacterial genomes from our initial set of 628 RR-like-containing organisms to conduct the simulations. From each of the 25 genomes, 2,000 pseudo-genomes were generated. Next, the RR-like neighboring genes (10 genes up and downstream) from each of the 50,000 simulations were inquired for the occurrence of R-M domains. The number of R-M domains in these regions were computed for each strain. For the vast majority of strains analyzed individually, the pseudo-genomes analysis showed that up to three R-M domain occurrences should be expected by chance in the RR-like gene neighborhoods (Figure 4A). Furthermore, we found that between five and seven R-M domains were found within in a maximum distance of 10 genes from the RR-like encoding gene in 41.9% of the 628 bacterial genomes analyzed. Whereas 9.9-15% of the 50,000 pseudo-genomes exhibited between five and seven R-M domains in the same genomic range. On the other hand, only 9.7% of the 628 genomes exhibit up to two R-M-related domains in the RR-like neighboring regions, whereas 49.3-56.8% exhibit the same frequency of R-M domains in the pseudo-genomes. These observations further underscored the conspicuous presence of R-M systems elements in the genomic contexts of the newly described RR-like orthologs. On top of these comparisons, statistical analyses showed that for all strains, the frequency of R-M domains in RR-like neighborhoods exhibited significant contrast (p*-*value < 0.0001) compared with the frequency found in the real data set (Figure 4B). The non-random genomic association of RR-Like and R-M-related genes strongly suggests a mechanistic relationship between these themes in a vast range of bacteria.

**Figure 4.**
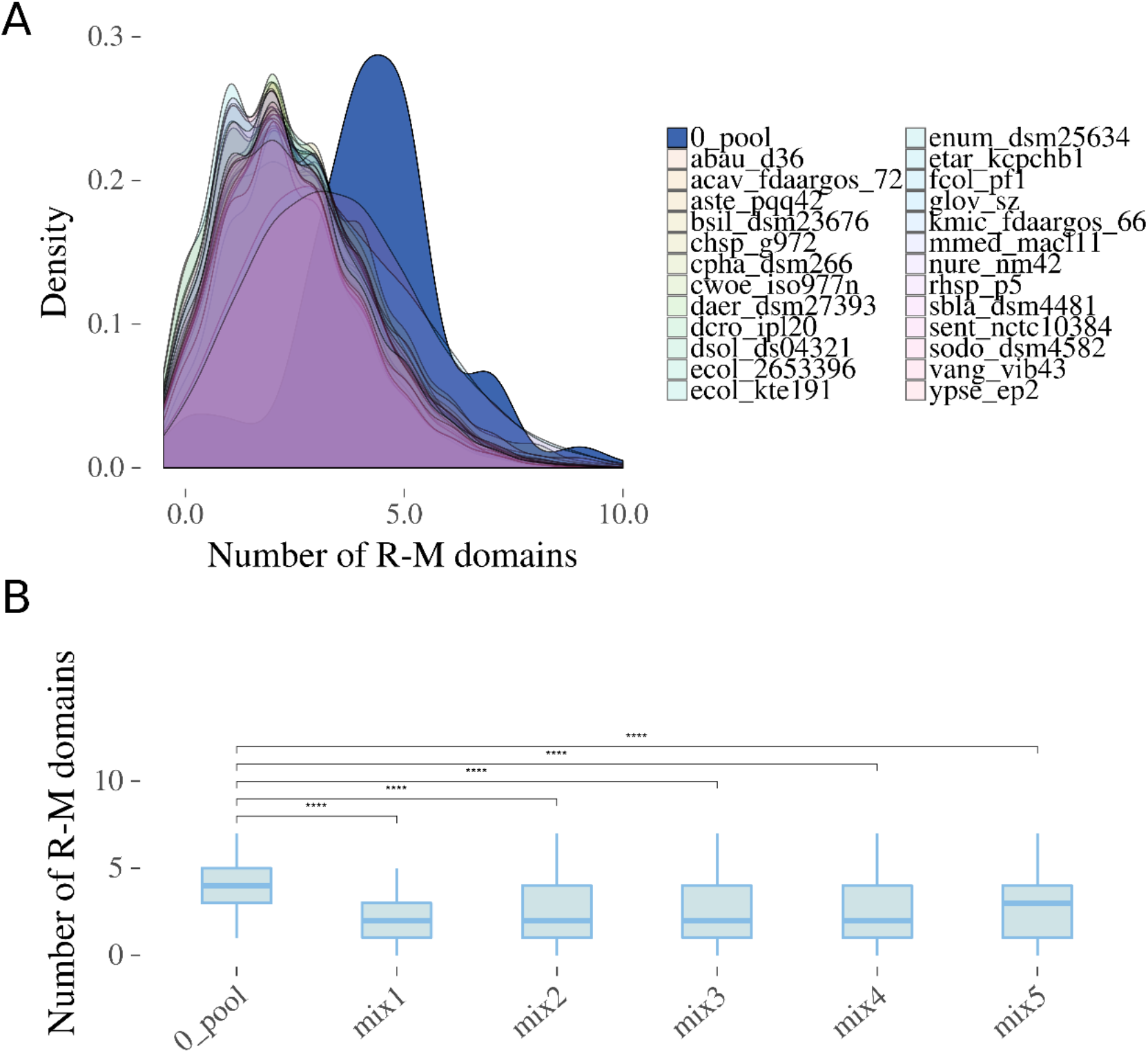
Distribution of R-M-related domains encoded in RR-like gene neighborhoods. **(A)** The density plot compares the distribution of genes encoding R-M domains found in close proximity to RR-like genes between the data set representing 628 actual bacterial genomes (0_pool) and the respective 2000 shuffled versions (pseudo-genomes) of 25 different genomes gleaned from the 0_pool. Species and strain names are abbreviated as follows: *Acinetobacter baumanni*i (abau_d36), *Aeromonas caviae* (acav_fdaargos_72), *Alteromonas stellipolaris* (aste_pqq42), *Brevibacterium siliguriense* (bsil_dsm23676), *Chryseobacterium* sp. G972 (chsp_g972), *Chlorobium phaeobacteroides* (cpha_dsm266), *Conexibacter woesei* (cwoe_iso977n), *Diaminobutyricimonas aerilata* (daer_dsm27393), *Devosia crocina* (dcro_ipl20), *Dickeya solani* (dsol_ds04321), *Escherichia coli* (ecol_2653396), *Escherichia coli* KTE191 (ecol_kte191), *Endozoicomonas numazuensis* (enum_dsm25634), *Edwardsiella tarda* (etar_kcpchb1), *Flavobacterium columnare* (fcol_pf1), *Geobacter lovleyi* (glov_sz), *Klebsiella michiganensis* (kmic_fdaargos_66), *Martelella mediterranea* (mmed_macl11), *Nitrosomonas ureae* (nure_nm42), *Rhodovulum* sp. P5 (rhsp_p5), *Shimwellia blattae* (sbla_dsm4481), *Salmonella enterica* subsp. *enterica* serovar Senftenberg (sent_nctc10384), *Serratia odorifera* (sodo_dsm4582), *Vibrio anguillarum* (vang_vib43), *Yersinia pseudotuberculosis* (ypse_ep2). **(B)** Box plots depicts the statistical analysis using one-tailed Student’s t-test between the 0_pool and five compilations (‘mix’) of the 25 pseudo-genomes previously simulated. Each mix contains five strains. T-test results are represented as follows: ns (p > 0.05), * (p <= 0.05), ** (p <= 0.01), *** (p <= 0.001), **** (p <= 0.0001). The mix datasets include the following above mentioned genomes: mix1 – abau_d36, acav_fdaargos, aste_pqq42, bsil_dsm23676, chsp_g972; mix2 – cwoe_iso977n, dear_dsm27393, dcro_ipl20, dsol_ds04321, ecol_2653396; mix3 – ecol_kte191, enum_dsm25634, etar_kcpchb1, fcol_pf1, glov_sz; mix4 – kmic_fdaargos, mmed_macl11, nure_nm42, rhsp_p5, sbla_dsm4481; mix5 – cpha_dsm266, sent_nctc10384, sodo_dsm4582, vang_vib43, ypse_ep2.

By further inspecting the RR-like regions (10 genes up- and downstream) with the REBASE annotation support, we found the prominent presence of type I and especially type II R-M system elements (Figure 5A). Although functional domains can often be coopted to perform their roles in different proteins from different systems (and can thus be classified into more than one R-M system type), the types I and II are still the most prevalent in these regions. Several nuclease domains from type II R-M systems were found in these vicinities, such as MvaI_BcnI (Pfam: PF15515), ParBc (Pfam: PF02195), EcoRII-C (Pfam: PF09019), ResIII (PF04851), HSDR (PF04313) and variants of the HNH domain (Figure 5B and TableS2). Specifically, the HNH_2 (Pfam: PF13391) is the best represented endonuclease domain in RR-like neighborhoods. The HNH_2 can be found in seven different genera of Gram-negative bacteria: Three from the Bacteroidetes (*Chryseobacterium, Flavobacterium*, and *Hymenobacter*) and four from the Proteobacteria (*Klebsiella, Pectobacterium, Salmonella*, and Yersinia) phyla (Figure 5B). Besides the type II-associated domains, the widespread presence of the Vsr endonuclease domain (Pfam: PF03852) was also observed. The Vsr-containing genes encode the very short patch repair (VSR) proteins. VSRs are often classified as V genes, and play a role in correcting mispairs in the DNA that arise following spontaneous deamination of 5-methylcitosine [35]. In this analysis we found Vsr-containing genes in 20 different genera, from four different phyla (Actinobacteria, Proteobacteria, Bacteroidetes, and Verrucomicrobia) encompassing: four γ-proteobacteria (*Acinetobacter, Azotobacter, Pseudomonas, Shewanella*), five α-proteobacteria (*Agrobacterium, Mesorhizobium, Pleomorphonas, Rhizobium, Sphingomonas*), two δ-proteobacteria (*Desulfovibrio, Geobacter*), five bacteroidias (*Bacteroides, Hallella, Hymenobacter, Porphyromonas, Prevotella*), one chlorobia (*Chlorobium*), one flavobacteriia (*Chryseobacterium*), one actinobacteria (*Diaminobutyricimonas*) and one verrucomicrobiae (*Akkermansia*). The most prevalent genomic organization of VSR-encoding genes across the different taxa was that in which VSR genes were adjacently located downstream to the RR-like genes (Figure 5B). These analyses underscored the solid association of RR-like with type II R-M systems, which may be preferentially recruited to work along with the newly discovered TCS-like system.

**Figure 5.**
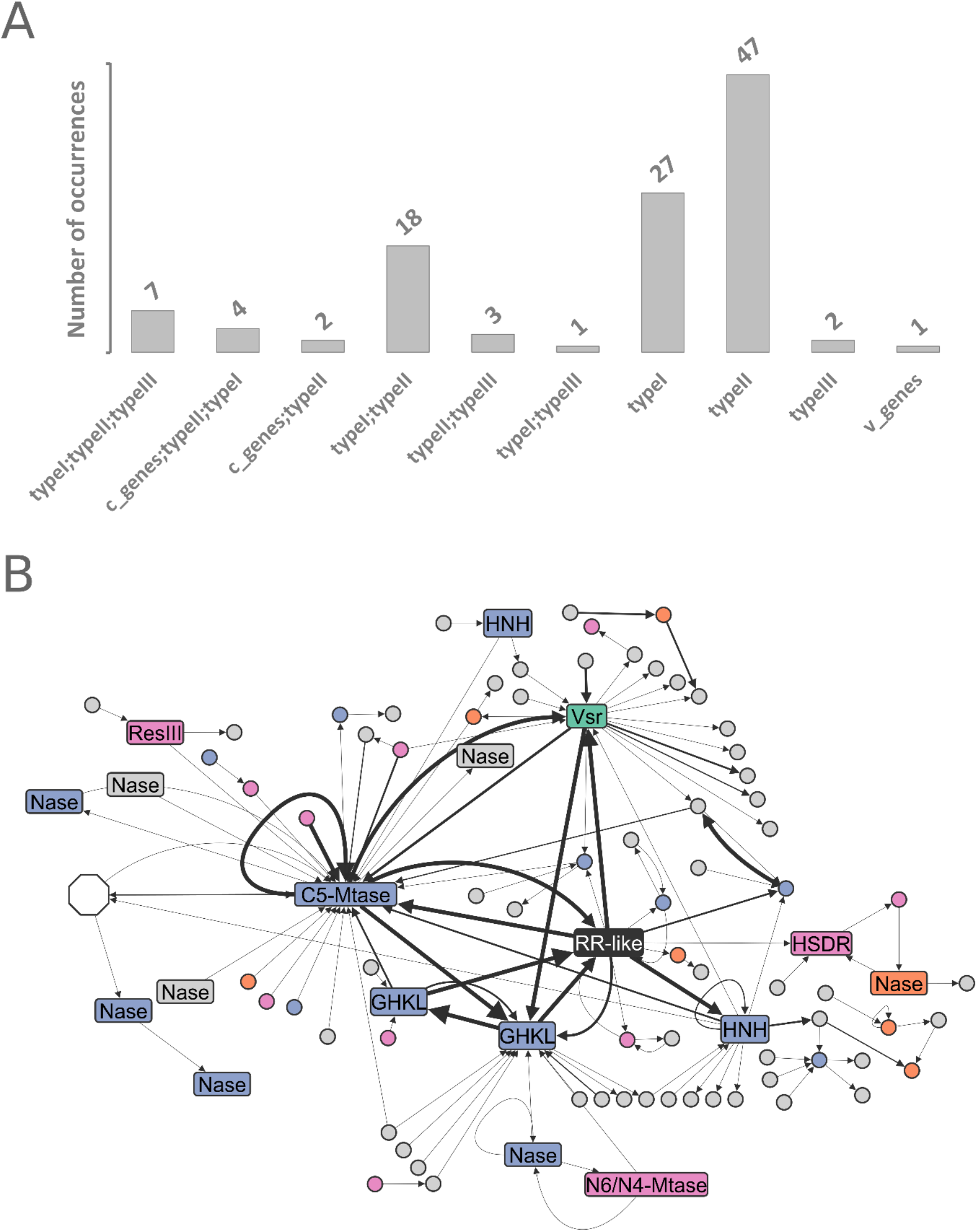
Classification of R-M systems identified in RR-like genomic regions. **(A)** Occurrence of distinct R-M types found in 10 genes up- and downstream of RR-like genes according to REBASE based on protein domains analysis. **(B)** Network representation using a subset (3 genes up- and downstream) of the analysis presented in (A) for improved visualization. The network shows the encoded domains with at least one R-M-related neighboring domain in the RR-like vicinities. Neighboring domains are linked with an arrow representing the order of appearance (5’ -> 3’) in the respective genomic regions. Line thickness is proportional to the number of genera in which a particular domain neighborhood was found. Different types of R-M systems according to REBASE database are highlighted in blue (type II), orange (type I), green (v genes), or pink (domains assigned to more than one R-M type). A white hexagonal shape represents 57 different non-R-M domains neighboring the C5-Mtase domain. Different gene families are labeled/abbreviated as follows: The Response_reg_2 encoding genes (RR-like), GHKL family domains, nuclease domains (Nase), HNH family domains (HNH), very short patch repair domains (Vsr), DNA_methylase domain encoded in C-5 cytosine-specific DNA methylase (C5-Mtase), N6_N4_Mtase domain encoded by N-6 adenine-specific DNA methylase or N-4 adenine-specific DNA methylase (N6/N4-Mtase), and ResIII or HSDR domains encoded by restriction enzymes.

### Phyletic distribution of protein architecture containing the newly characterized HK-like

The next step was to consolidate the newly found domain profiles based on the representative sequences respectively from OG_4 and OG_5. Consolidation of the new profiles enabled us to perform broad domain-oriented searches of Response_reg_2 and HEF_HK conserved motifs against large databases using hmmsearch [36]. By conducting this sensitive search, we aimed to test the results from the previous analysis and assess the prevalence of the new domains across the bacterial lineages.

Hence, Ensembl, UniprotKB and Reference proteomes (uniprotrefprot) were inspected for the presence of the two domains, which returned respectively from these databases: 529, 163, and 50 hits for Response_reg_2 and 199, 139 and 52 hits for HEF_HK. As expected, based on our first analysis, the Response_reg_2 domains were detected in proteins lacking any other currently known domains. On the other hand, the HEF_HK proteins exhibited diversity in terms of associated known domains. These sequences contained GHKL superfamily motifs (i.e. HATPase_c, and HATPase_c_3), which strikingly confirmed the pattern observed in the first section of this study. Furthermore, the variable presence of functional domains could be exploited in order to glean additional information on the phyletic distribution of the HEF_HK-containing proteins. In order to achieve that, all HEF_HK significant hits found by hmmsearch in the three databases mentioned above were acquired, and then merged with the OG_5 sequences from the previous analyses. Next, this merged data set was filtered to remove redundant sequences, which typically belong to different strains of the same species. This step prevented inflated representation of species that were prevalent in public databases. This resulted in a data set including only unique sequences to be further analyzed. Next, aiming to ensure that only bona fide HEF_HK-containing proteins were to be included, we performed an additional in-house domain profile recognition on the unique sequences set with a stringent setting (e-value: 1e-05) using hmmscan (TableS3). The results from the 222 unique bona fide HEF_HK-containing sequences showed that most of these were found in organisms from Proteobacteria phylum (69.8%). Furthermore, it underscored the remarkable association of this domain with two other domains: (a) the HATPase_c_3, found within N-terminal regions (83.3%), and/or the (b) HATPase_c within C-terminal regions (93.7%) [TableS3]. In this context, the predominant domain organization observed is the one including HATPase_c_3+HEF_HK+HATPase_c (group 1), represented in 174 out of 222 sequences (Figure 6A). Followed by group 2 (34/222), which have lost the N-terminal HATPase_c_3 (i.e. HEF_HK+HATPase_c). Group 3 (11/222), instead, comprised those architectures in which the C-terminal HATPase_c was lost (i.e. HATPase_c_3+HEF_HK). As the most unusual architecture, group 4 (3/222) conserved only the HEF_HK, with no other known domain.

**Figure 6.**
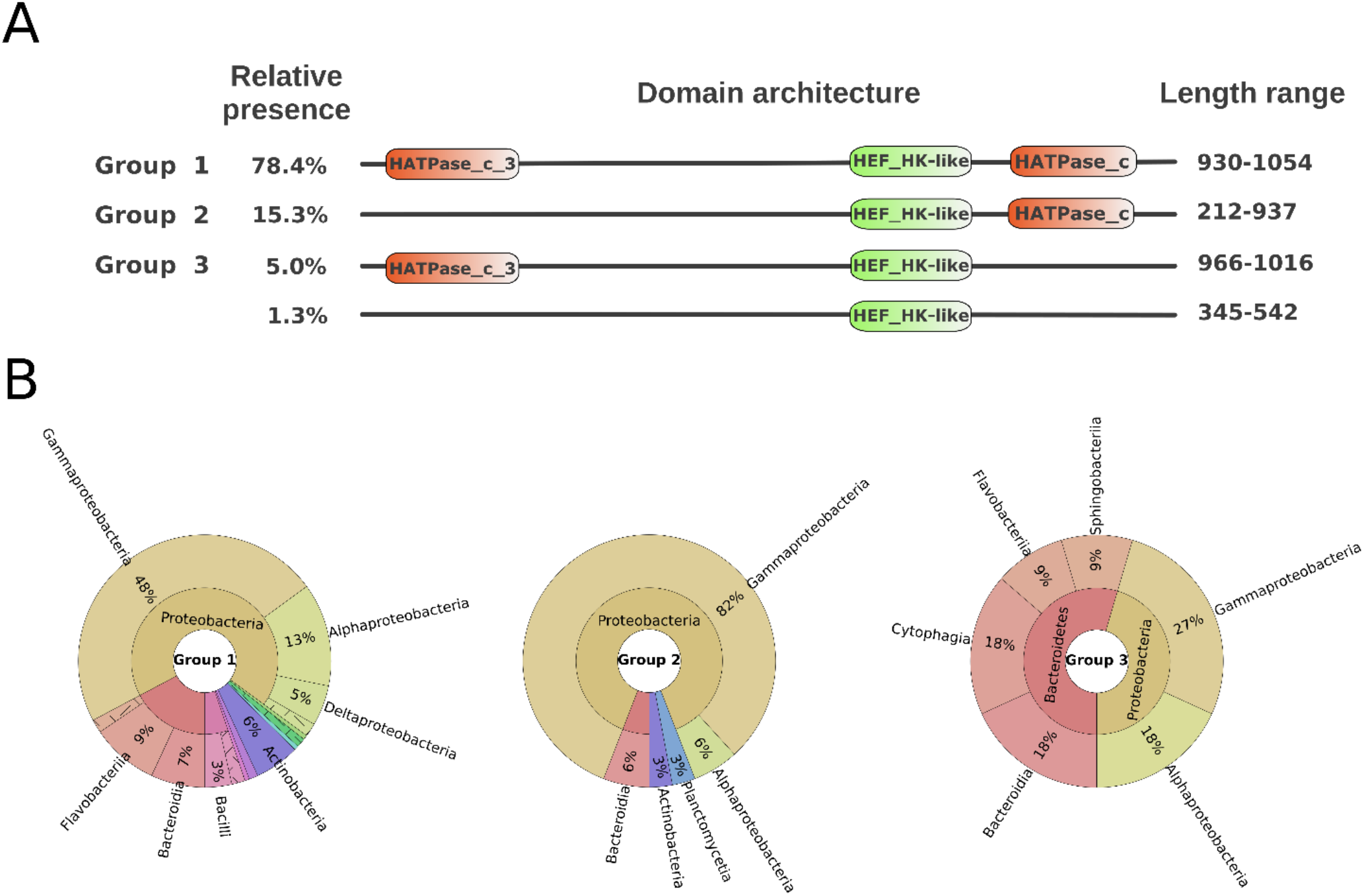
HK-like protein domain architectures presence in bacterial lineages. **(A)** Different architectures found in the 222 unique bona fide HEF_HK-containing sequences retrieved from Ensembl, UniprotKB and Reference proteomes (uniprotrefprot). The three prevalent architectures are labeled as Group 1, 2 and 3 and their relative presence in the entire set of sequences is adjacent to the group labels. Domains recovered from the Pfam database are represented in orange, whereas the newly consolidated HEF_HK appears in green. Longest and shortest sequence lengths of each group are depicted adjacent to each architecture. **(B)** The phyletic distribution of the three prevalent architecture groups is depicted by composite pie charts.

Notably, group 1 was the only architecture found in HEF_HK-containing sequences conserved in Betaproteobacteria, Deltaproteobacteria or in the Firmicutes phylum (Figure 6B and TableS3). Numerous orders of the Alphaproteobacteria class were shown to conserve any of the three major groups (i.e. 1, 2 and 3). Among these orders, Rhodobacterales seemed to obligatorily conserve the C-terminal signature (HATPase_c), compared to the N-terminal HATPase_c_3 may not be essential. Contrarily, in the Rhizobiales order, the N-terminal HATPase_c_3 seem to be mandatory. The Gammaproteobacteria class encompasses most of the organisms carrying HEF_HK-containing sequences (51.8%). Within this class, bacteria from Yersiniaceae (including *Yersinia* and *Serratia* spp.) and Pectobacteriaceae (including *Pectobacterium* and *Dickeya* spp.) families are constricted to group 1. Conversely, Moraxellaceae and Vibrionaceae families may conserve either group 1 or 2, meaning that the C-terminal HATPase_c is essential in these organisms ‘sequences. These results underscore the wide diversity of organisms in which the HEF_HK domain is conserved, and the contrasting conservation for the companion HATPase domains in different phyletic groups.

## DISCUSSION

In this study, two novel conserved domains were characterized: HEF_HK and Response_reg_2. The newly described domains conserve characteristic secondary structure organization canonically described DHp and REC domains found respectively in HKs and RRs. In HK proteins, although several DHp domain variants have been described, some key features must remain unaltered such as the structure of the kinase core and the adjacent nucleotide binding fold [37]. The archetypal organization among the highly conserved domains of HK proteins includes five motifs known as H, N, G1, F and G2 boxes [38]. The H box harbors the core His residue and remains typically conserved adjacently to the C-terminal nucleotide-binding pocket which encompasses the N, G1, F and G2 boxes [39]. As a highly conserved structure, the C-terminal CA domain containing the nucleotide-binding pocket could be promptly detected in the sequences from our acquired dataset by exhibiting the HATPase signature. Within these sequences, the structure reminiscent of an H box could be detected adjacently to the C-terminal HATPase_c domain (Figure 6A). Of the sequences carrying a HEF_HK domain, 83.4% also conserved an N-terminal HATPase_c_3 (Pfam: PF13589) domain. The HATPase_c_3 domain is classified into the GHKL superfamily similarly to HATPase_c (Pfam: CL0025). By looking at the complete set of sequences carrying the HATPase_c_3 domains, the strikingly strong preference for N-terminal localization was clear, contrarily to the C-terminal preference observed for HATPase_c domain [32]. The N-terminal localization of HK-associated HATPase_c_3 occupies the classic site of sensor domains in HK proteins [40]. Intriguingly, none of the canonical DHp domains can be found in HATPase_c_3 containing proteins [32]. These observations could suggest a specific recruitment of HATPase_c_3 domains to play a role as sensor in the newly predicted HEF_HK containing HKs. Since the HEF_HK containing proteins are predominantly soluble (Figure 2D), it is possible to speculate that HATPase_c_3+ HEF_HK containing proteins could be involved in interactions with intracellular concentrations of <u>n</u>ucleoside triphosphate <u>m</u>olecules (NTP).

Characterization of TCSs utilizing sequence analysis and extensive comparative genomics has been largely exploited in the last decades, comprising a powerful strategy for identification of new systems [41-43]. Besides revealing the solid linkage between genes encoding HEF_HK- and Response_reg_2-containing products, this approach also shed light on the striking conservation of upstream genes to this duplet encoding methyltransferases [TableS2]. The DNA_methylase domains integrate the <u>DN</u>A <u>m</u>ethyltransferases (DNMT) protein family which became specialized in cytosine-specific methylation [44]. In this context, DNA methylation in prokaryotes has been largely associated with restriction-<u>m</u>odification (R-M) systems, which typically contain by two opposing utilities: endonucleases (performs double-strand DNA cleavage) and DNA methyltransferase (prevents double-strand cleavage by the cognate endonuclease) [45]. The linkage between R-M systems and predicted ATPase-encoding genes has been previously observed by [46], however the presence of a predicted response regulator has not been revealed thus far.

The modification systems are effective barriers against bacterial lineages of varied descent which contain different epigenetic identities as defined by their own R-M systems [47]. Since some R-M systems can occasionally target the host genome if eliminated, they are typically referred to as selfish elements as they promote their own survival in the cell lineages [47-49]. Thus, if the concentration of M proteins is not sufficient to protect the host DNA against cognate R endonucleases in the cell, this may cause cell death [50]. The overrepresented presence of R-M-related domains in the vicinity of HEF_HK-/Response_reg_2-containing genes observed in a wide range of bacterial genomes corroborates that the association between these themes does not occur by chance (Figure 5 and TableS2). Hence, one might speculate that these newly discovered domains could be involved in signaling cascades controlling, directly or indirectly, restriction-modification systems.

The N-terminal presence of the GHKL domain HATPase_c_3 in HEF_HK-containing proteins is a well-established characteristic feature of the Microrchidia (MORC) ATPase family, originally described as a necessary gene for the completion of mammalian spermatogenesis [51]. Subsequent investigations revealed the presence of MORC family members in prokaryotes [52], as well as in the major crown group lineages of eukaryotes [53,54]. Although mechanistic details concerning the majority of MORC’s functions remain elusive, they constitute a widespread family primarily involved in diverse processes associated with epigenetic regulation [55]. In *Arabidopsis thaliana*, for example, AtMORC1 and AtMORC6 have been implicated in gene silencing through chromatin condensation [56]. Another MORC1 (SlMORC1 from *Solanum lycopersicum*) was further reported to exhibit similarities with type II topoisomerases [57]. Furthermore, the ability to form dimers through the N-terminal GHKL domain was elucidated in the murine MORC3 protein [58]. In contrast to the recent advances in the understanding of eukaryotic MORCs, the molecular and biochemical aspects of the bacterial MORCs remain poorly understood. Hence, it is currently unclear whether the focus in chromatin superstructure manipulation is conserved across the eukaryotic and prokaryotic members. In prokaryotes, the MORCs were classified as part of a larger radiation that includes a number of GHKL proteins termed paraMORCs [52]. The paraMORCs were shown to remain preferably in genomic regions co-inhabited by R-M systems, endonucleases and helicases [52].

The striking similarity of sequence and contextual signatures between the paraMORCs and HEF_HK-encoding genes strongly suggests that this newly found HK-like family may have branched from the paraMORCs through the acquisition of the C-terminal HATPase_c and HEF_HK domains. Altogether, this study revealed two novel protein conserved domains and shed light to the possibility of a TCS-like system undertaking a regulatory role mechanistically linked to R-M systems in a large variety of bacterial lineages.

## MATERIAL AND METHODS

### Sequence search, orthology and gene-neighborhood screenings

The initial search for RR-similar sequences was conducted by using TBLASTN 2.8.0+ [59] in 5 iterations on RefSeq Genomes database available on NCBI webpage (https://www.ncbi.nlm.nih.gov), allowing up to 1000 positive hits restricted to Bacteria (taxid:2). The query used for this search is 600 aa long and was obtained from *Pectobacterium carotovorum* subsp. *brasiliense* (KS44_RS10365). A total of 684 positive hits returned, for which 628 came from bacterial strains with support of genome-wide information. These datasets were then obtained, enabling subsequent gene-neighborhood screenings in the strains in which the hits were found. Full records in Genbank format of the 628 entries representing their respective genomic constructs (complete genome, scaffold, or contig) containing the KS44_RS10365-similar sequences were downloaded from RefSeq database. Additional ecological and taxonomical information on these 628 strains were then gleaned from different sources, including NCBI taxonomy-, Uniprot-, and PATRIC bacterial databases [60-62]. Taxonomic records of interest include the bacterial class, order and family.

Following the preliminary sequence search, two distinct methods will be carried out aiming to provide relevant information on the obtained sequences for subsequent gene-neighborhood screening. First, the OrthoMCL [30] pipeline was applied in order to predict homology relationships among the sequences by orthologous clustering based on MCL (inflation = 1.5). Each <u>o</u>rthologous <u>g</u>roup (OG) is numerically labelled for easy identification (e.g. OG_1, OG_2, etc.). Second, all sequences were scanned for known conserved domains using HMMER3 package [31] supported by the PFAM domains database [32]. In order to combine both sources of information in the gene-neighborhood screenings, the genomic coordinates were then integrated to these results by custom PERL scripts (https://www.perl.org). The identification of R-M-related domains in this analysis was achieved by retrieving the gold standard sequences from REBASE database [34] and performing in-house detection of functional domains using HMMER just as described above. This information was integrated into the gene-neighborhood screenings, and the resulting network was then generated by using Cytoscape software [63].

### Sequence alignment, phylogeny and domain analysis

The orthologous sequences of both OG_4 (RRs) and OG_5 (HKs) were aligned by Clustal Omega [64] and PROMALS3D [65], and visualized by Jalview [66] in comparison with canonical REC and DHp domains obtained from Pfam database (https://pfam.xfam.org/). All alignments performed involving canonical REC and DHp domains use the Pfam ‘seed ‘entries available in the online repository for each described domain, which includes only representative sequences carrying the respective domains. This approach aimed to comparatively assess the conservation of key RR and HK residues in the sequences from OG_2 and OG_4. Secondary structures were also inferred in the alignments by using JPred4 [67], and membrane topology predicted by TopCons server [68]. To establish comparison with OG_2 sequences, we obtained the domain families from Pfam clan CL0304 carrying a highly conserved Asp residue (PFAM: PF00072 - Response_reg; PF16359 - RcsD_ABL; PF09456 – RcsC) to perform alignments with OG_2 sequences. Aside from Response_reg domain, the other combined alignments with OG_2 sequences showed poor quality and were hence discarded. Sequences from OG_4 were aligned with domains families from Pfam Clan CL0025 carrying highly conserved His residue (PFAM: PF00512 - HiskA; PF07568 – HiskA_2; PF07730 - HiskA_3; PF07536 – HWE_HK), resulting in a good quality combined alignment. Individual alignments including only sequences carrying Response_reg_2 and HEF_HK domains respectively were then performed in order to determine the limits of each domain. These limits were determined based on secondary structure analyses performed using JPred. The Response_reg_2 and HEF_HK domains were then cut from the alignments, and filtered to avoid sequence redundancy, and then supplied to SMS [69] for evolutionary model selection. Next, the domain sequences were supplied to FastTree [70] in order to reconstruct the domains phylogeny. The cut domains from Response_reg_2 and HEF_HK were converted into HMM-profiles, which were consolidated by *hmmbuild* and *hmmpress* from HMMER3 package. After the domain profiles were consolidated (Response_reg_2 and HEF_HK), these were used in a high sensitivity search in public databases through hmmersearch under default parameters [36]. The positive matches were obtained, merged with the sequences from the respective orthologous groups, and then filtered to avoid sequence redundancies. The non-redundant sets of Response_reg_2 and HEF_HK were then subjected to in-house domain scanning under stringent criteria (e-value < 1e-05) using hmmscan, in order to avoid low-confidence domain prediction. The 3D model predictions of the conserved motifs in RR-like and HK-like were carried out by using RaptorX [71] which selects the most suitable PDB structure for each analyzed sequence and uses it as reference. Four randomly picked representative sequences from both the RR-like (BJP29_RS22595 – PDB ID: 2ZWM; H147_RS0115650 – PDB ID: 1A2O; B038_RS0119475 PDB ID: 3Q9S; VOL_RS02805 PDB ID: 3Q9S) and the HK-like (CFY84_RS11065 – PDB ID: 5OO7; HMPREF0127_RS22520 – PDB ID: 5OO7; WKA_RS00445 – PDB ID: 5OO7; KORLAC_RS15945 – PDB ID: 5JRW) groups were analyzed using this method. Subsequent visualization, image rendering of domain models used Pymol [72], in combination with EnvZ (PDB ID: 5B1N) and CheY (PDB ID: 2FKA) obtained from PDB online repository [73] for comparison.

## AUTHOR CONTRIBUTIONS

Conceptualization, D.B-R. and L.N.M.; methodology, D.B-R., W.J.S.P., and L.N.M.; formal analysis, D.B-R., and W.J.S.P.; writing—original draft preparation, D.B-R. and W.J.S.P.; writing—review and editing, D.B-R and L.N.M.; funding acquisition, L.N.M. All authors have read and agreed to the published version of the manuscript.

## FUNDING

This research was funded by the National Research Foundation (NRF), South Africa through Competitive Funding for Rated Researchers (CFRR) 98993. W.J.S.P. studentship was supported by The Research Technology Fund (RTF) 98654. D.B-R. was supported by The University of Pretoria Post-Doctoral Fellowship. Any findings and/or recommendations expressed here are those of the author(s) and the NRF does not accept any liability in this regard.

## ACKNOWLEDGEMENTS

We would like to acknowledge the expert assistance of Dr Divine Y. Shyntum with insights on microbial genetics and genomics in the early stages of this work.

## CONFLICT OF INTEREST

The authors declare no conflicts of interest.

## DATA AVAILABILITY

The protein domain profiles described in this study are available online on the PFAM database and can be found respectively on these addresses: https://xfamsvn.ebi.ac.uk/svn/pfam/trunk/Data/Families/PF19191/ (HEF_HK) and https://xfamsvn.ebi.ac.uk/svn/pfam/trunk/Data/Families/PF19192/ (Response_reg_2).

## Notes

### Competing Interest Statement

The authors have declared no competing interest.

